# A G2 Checkpoint Arrests *Cryptococcus neoformans* Cell Division in response to Hypoxia

**DOI:** 10.64898/2026.06.30.735586

**Authors:** Huarui Zhou, Claudia A. Petrucco, Ashley H. Lim, Steven B. Haase

## Abstract

Saturated cultures of the pathogenic yeast, *Cryptococcus neoformans*, arrest as unbudded cells in the G2 phase of the cell cycle. As cells divided and cultures saturated, we found that oxygen levels in the culture medium dropped nearly tenfold. When saturation-arrested cultures were re-oxygenated without adding fresh growth medium, cells immediately formed a bud and then underwent mitosis. Thus, the arrest is due to low oxygen concentration rather than nutrient depletion. Because the G2 arrest was associated with unbudded cells, we asked whether *C. neoformans* cells have a morphogenesis checkpoint that blocks mitosis until cells can form a bud. Inhibition of budding by treatment with Latrunculin A also led to G2 arrest, and we determined that this arrest is dependent on the CDK inhibitory kinase, Swe1. This finding suggests that *C. neoformans* possesses a morphogenesis checkpoint analogous to that in the distantly related *Saccharomyces cerevisiae*. We also demonstrated that Swe1 is required to enforce the hypoxia-induced G2 arrest. We propose that hypoxia inhibits budding in *C. neoformans*, which in turn triggers a morphogenesis checkpoint to arrest cells in G2 even when nutrients are plentiful.

## INTRODUCTION

In liquid culture, microorganisms such as bacteria and yeast typically transition through a predictable set of growth phases. Immediately after inoculation into fresh media, a lag phase in which cells exit quiescence is followed by a log phase where cells undergo exponential growth. As cell density increases and nutrients become limiting, cells enter a stationary phase associated with cell-cycle arrest. This pattern has been extensively studied in *Escherichia coli* ^1^ and the model budding yeast *Saccharomyces cerevisiae* ^2^. *C. neoformans* is widely thought to follow a similar pattern, entering stationary phase as nutrients are depleted^3–5^.

The stationary phase of *C. neoformans*, is characterized by increased cell size and unbudded G2 arrest^3^, which has recently been linked to several pathogenic processes. First, entry into the stationary phase has been shown to promote persistence against antifungal drugs such as amphotericin B^5^. Additionally, G2-unbudded cells in the stationary phase have been identified as precursors of Titan cells^4^, which are polyploid, enlarged cells that significantly enhance virulence, stress adaptation, and resistance to the host immune system^6–9^. Therefore, investigating the stationary phase of *C. neoformans* is crucial for understanding *Cryptococcal* infections.

Entry into stationary phase is an active decision that cells make in response to dwindling nutritional resources rather than simply stopping proliferation due to starvation. In fact, pathways that enable this decision help protect cells from starvation and loss of viability by forcing cells to exit the cell-division cycle into a quiescent phase before nutrients are fully depleted^10,11^. Similar intracellular signaling pathways, called checkpoints, direct cells to arrest when they detect perturbations of cell-cycle events. For example, the DNA damage checkpoint monitors DNA damage and blocks cell-cycle progression until the damage can be repaired. In this case, the ability to pause the cell cycle maintains genome stability and cell viability in the face of DNA damage^12,13^. Additional checkpoints include (but are not limited to) the DNA replication checkpoint, which prevents entry into mitosis until DNA replication is complete, and the spindle assembly checkpoint, which prevents the metaphase/anaphase transition until the chromosomes are all properly attached to the mitotic spindle^14,15^. A key feature of these pathways is that they are reversible, so that when damage is repaired or events are completed, the cells re-enter the cell cycle and continue proliferation.

*C. neoformans* cells grown to saturation in liquid culture were arrested as unbudded cells in the G2 phase rather than unbudded cells in the G1 phase with unreplicated DNA, as generally observed for stationary phase cells^10^. We found that surprisingly, these cells were not starved for nutrients, but instead were arrested due to low oxygen levels (hypoxia). Cells responded to hypoxia by activating a checkpoint that blocked mitosis, resulting in a G2 arrest. The checkpoint appears to be related to hypoxia-induced inhibition of budding, as we uncovered evidence that *C. neoformans* has a budding/morphogenesis checkpoint like that found in *S. cerevisiae* ^16^.

## RESULTS

### Nutrient depletion is not the cause of the saturation arrest

To understand why *C. neoformans* cells arrest at saturation, we analyzed DNA content, budding index, and cell density in a population of *C. neoformans* cells growing in YEPD (1% yeast extract, 2% peptone, 2% dextrose) medium as they moved from lag phase to exponential growth to saturation arrest. Consistent with previous studies^3,5^, cells were arrested as unbudded cells in the G2 phase after reaching saturation (Figure 1). However, several features of this arrest seemed inconsistent with a simple nutrient depletion response.

**Figure 1:**
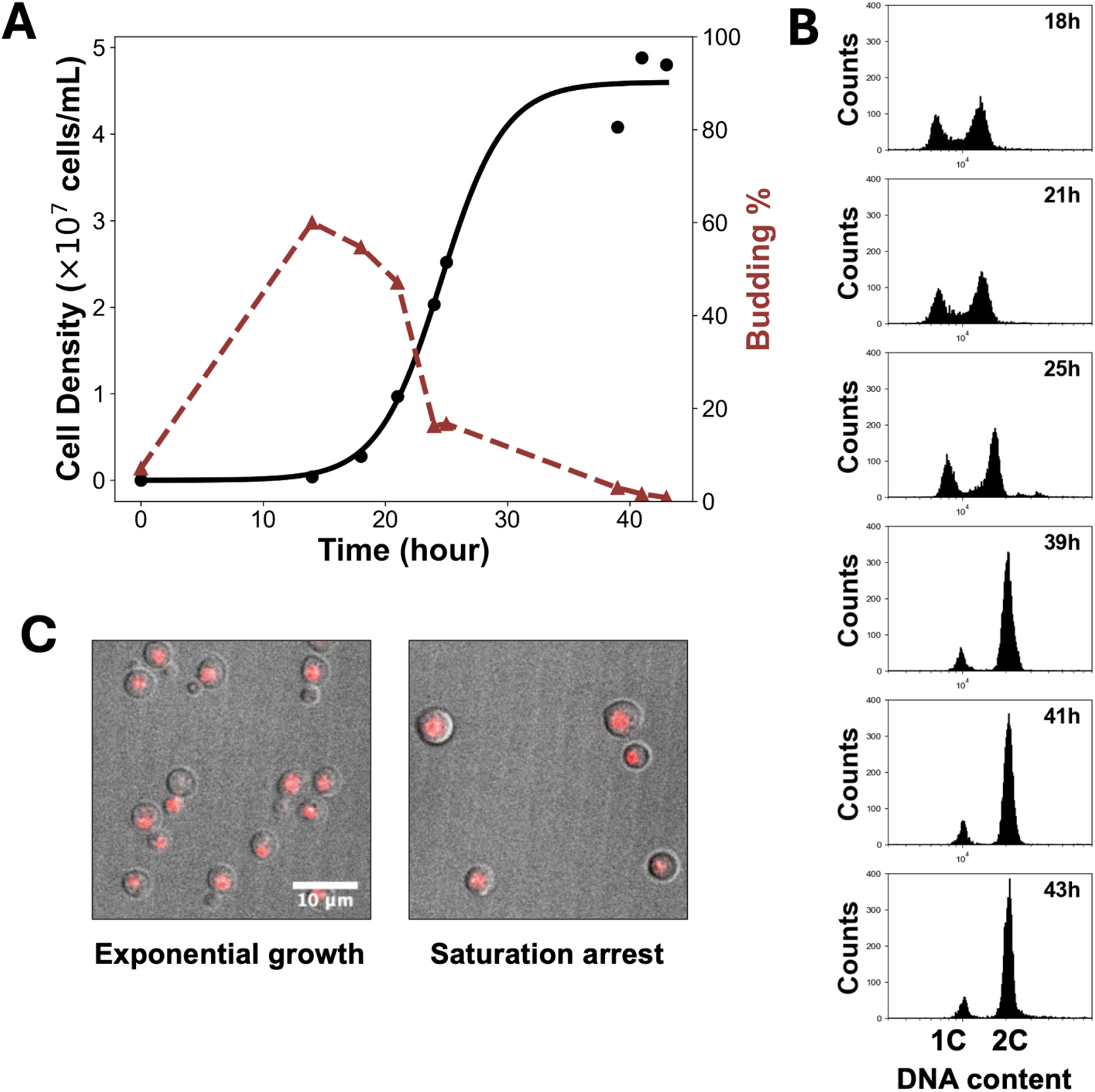
*C. neoformans* H99 cells arrest unbudded in the G2 phase at saturation. (**A**) Growth curve and budding index of *C. neoformans* H99 grown in YEPD. The growth curve was fitted using a logistic model. (**B**) DNA content of the cells collected in (**A**), stained with SYTOX Orange (**C**). Representative images of cells during exponential growth and at saturation arrest. DNA was stained with SYTOX Orange.

First, *S. cerevisiae* cells arrest at high density as unbudded cells in G1 phase^10^. Mammalian cells also arrest in G1 in response to being starved for serum growth factors^17,18^. Second, when we released these saturation-arrested *C. neoformans* cells into fresh YEPD media, they budded immediately, producing a population of cells that moved very synchronously through successive cell cycles as observed previously^19^ (Figure S1A). The synchrony is also observed in another *Cryptococcus* species, *C. gattii* (Figure S1B). The synchrony generated by release from saturation is comparable to that induced by alpha-factor in *S. cerevisiae* (Figure S1C), and contrasts with that of *S. cerevisiae* under nutrient-depleted conditions, which typically do not exhibit strong synchrony upon release from quiescence^2^. Third, we also observed that the percentage of budded cells begins to drop just as the cells enter exponential growth around 13 hours (Figure 1A) rather than dropping as the cells enter the saturation arrest. The drop in budding percentage in the log phase does not reflect a failure to bud, as cells would not be able to divide rapidly without producing a bud. Consistent with previous observations^20^, the drop in budding percentage is likely due to a shift of bud emergence from early in the cell cycle to later in the cell cycle, near mitosis/cytokinesis, which shortens the window during which cells are budded. Eventually, as cells reach saturation arrest, bud emergence stops entirely. Taken together, these observations suggested that cells might have altered the cell cycle in response to some signal rather than the depletion of nutrients.

To directly test whether nutrient depletion was responsible for the arrest, we pelleted cells from a saturation-arrested culture and then resuspended them in the spent medium from the arrested culture (conditioned medium) at lower density. If nutrients were exhausted, cells should remain arrested in the conditioned medium. However, we observed that the arrested cells immediately resumed budding and re-entered exponential growth (Figure 2 A, B), indicating that the conditioned medium still contained sufficient nutrients to support cell growth and division. Therefore, nutrient depletion is not the cause of the saturation arrest.

**Figure 2:**
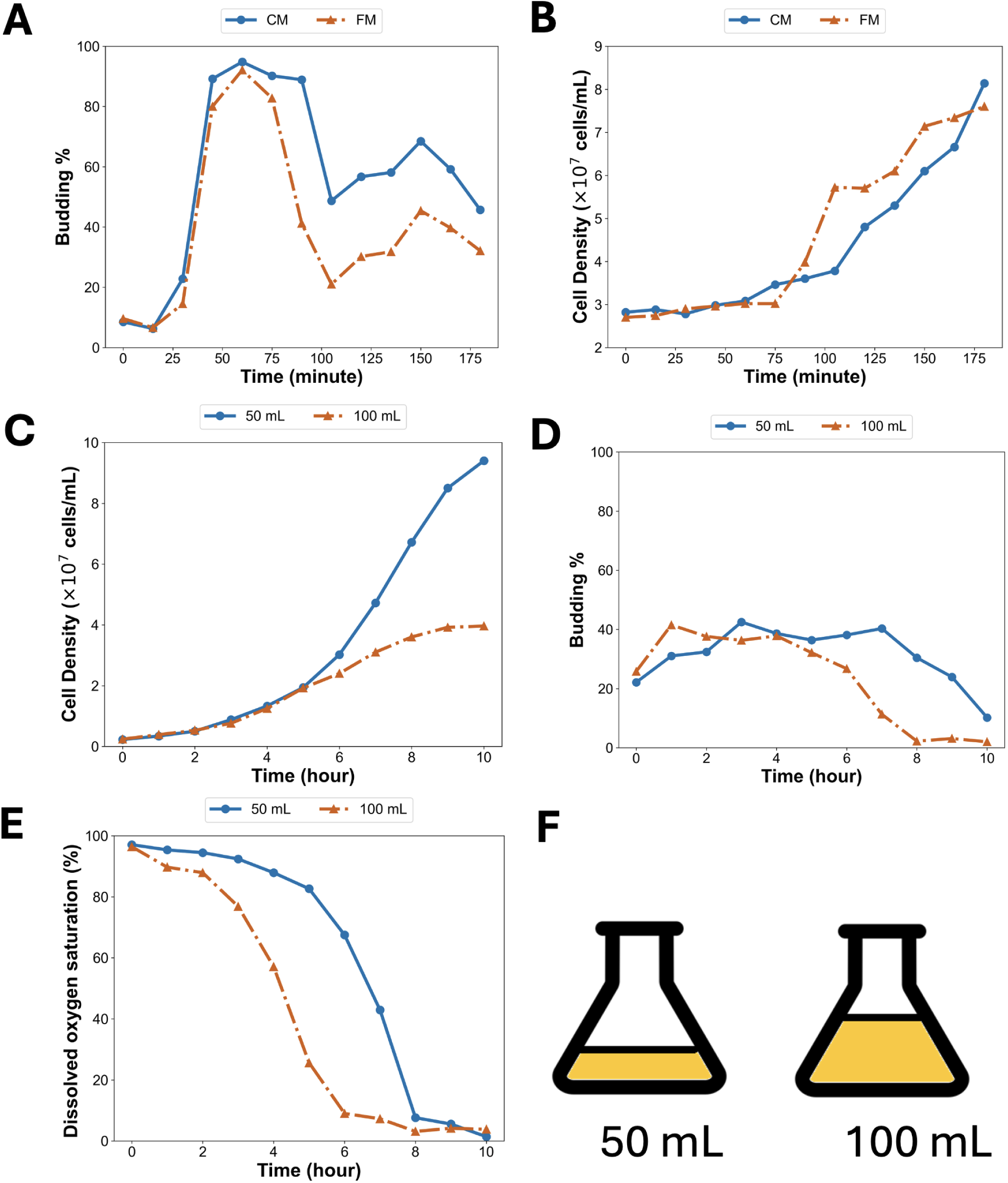
Nutrient depletion is not the cause of the saturation arrest. Budding index (**A**) and growth curve (**B**) of saturation-arrested H99 cells resuspended into conditioned medium (CM) or fresh medium (FM). Growth curve (**C**), budding index (**D**), and oxygen level (**E**) of H99 cells grown in 50 mL or 100 mL of medium. (**F**): Diagram showing how culture volume affects surface area and headspace, thereby affecting oxygen availability.

### Hypoxia Induces G2 Arrest

We then asked what triggered the saturation arrest. We had observed that the cell density at saturation varied depending on the amount of growth medium in a particular sized flask (approximately 5 × 10^7^ cells/mL in 100 mL media and 1.5 × 10^8^ cells/mL in 50 mL media) (Figure 2 C). This observation hinted that oxygen availability might play a role, as the 50 mL flask has more surface area for oxygen exchange (Figure 2 F). To test this idea, we monitored oxygen levels during cell growth in different volumes of growth media and found that dissolved oxygen levels dropped faster in 100 mL of medium as compared to 50 mL culture (Figure 2 E). Regardless of the volume of the medium, cells stopped budding and arrested in G2 when dissolved oxygen dropped below ten percent, suggesting that the 100 mL culture was arresting earlier due to a more rapid drop in dissolved oxygen.

Based on this observation, we hypothesized that hypoxia induced the G2 arrest at saturation density. To further test this hypothesis, we re-infused saturated cultures with oxygen by continuously bubbling air into the culture medium. As a result, oxygen levels rose rapidly, and the saturation-arrested cells immediately resumed budding and dividing (Figure 3 A, B, C, and Figure S2 A). We performed the same air-infusion treatment on two additional *Cryptococcus* species, *C. gattii* R265 and *C. deneoformans* JEC21. Both species responded similarly, resuming budding immediately upon air infusion (Figure S3 A, B, C), suggesting that hypoxia-induced saturation arrest is conserved across *Cryptococcus* species.

**Figure 3:**
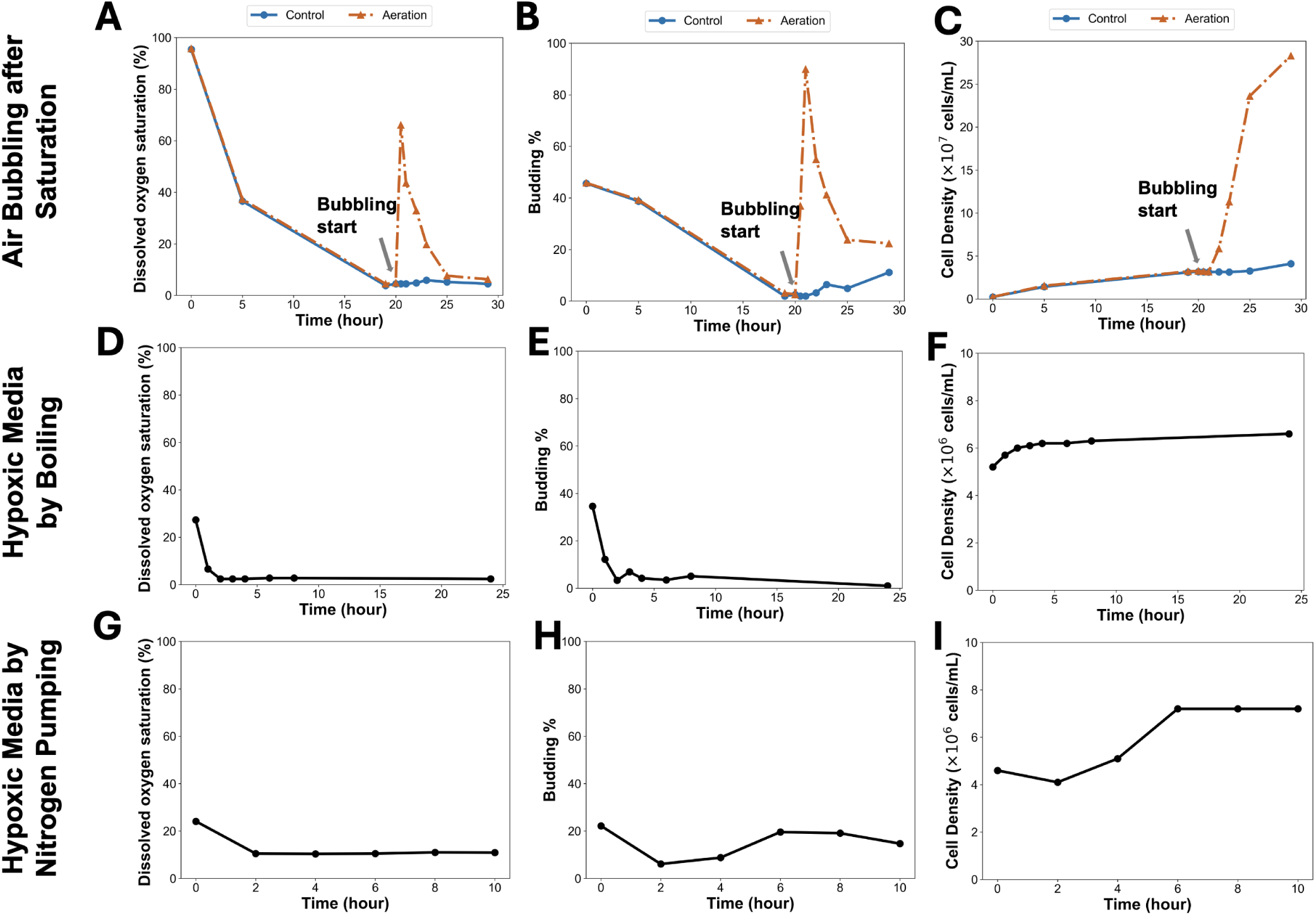
Hypoxia Induces G2 Arrest. Oxygen level, budding index, and growth curve of H99 cells without air bubbling (control) or with air bubbling (**A–C**), and of cells transferred to hypoxic medium generated by boiling (**D–F**) or by nitrogen pumping (**G–I**). Air bubbling was achieved by continuously pumping fresh air into the medium after the time indicated by the arrow.

The observations described above could be consistent either with the loss of O_2_ or the accumulation of CO_2_. *Cryptococcus* is known to be sensitive to CO_2_ levels^21,22^, so to distinguish between O_2_ depletion and CO_2_ accumulation, we produced an acute hypoxia condition using two methods. First, we boiled the YEPD medium to remove all dissolved gases, and then inoculated cells into the degassed medium after cooling. Under this condition, cells stopped growing before completing a single division cycle, and the majority arrested in G2 phase (Figure 3 D, E, F, and Figure S2 B). Using an independent method for depleting O_2_ and CO_2_ from the growth medium, we bubbled N_2_ into the medium to remove other gases, and observed a similar rapid arrest (Figure 3 H, I, and J). Together, these observations indicate that hypoxia rather than CO_2_ accumulation induces the G2 arrest in saturated culture.

### The G2 arrest may be triggered by a checkpoint responding to the inhibition of budding

In response to severe hypoxia at saturation, we observed both an inhibition of budding and a G2 arrest of the cell cycle. In principle, it could be that inhibition of budding causes G2 arrest, or that G2 arrest blocks budding, or that the budding and G2 arrest are both independent effects of hypoxia. To examine the relationship between the two observations, we used live-cell imaging on a culture of cells released from the saturation into fresh, oxygenated medium. When *C. neoformans* cells with histone H4-GFP^23^ were released from hypoxic arrest, bud emergence and growth always preceded mitosis (Figure 4 and Figure S4). Consistent with previous observations^23^, the nuclei move into the bud before anaphase begins (Figure 4 and Video S1), suggesting that nuclear movement into a bud is required for the initiation of anaphase. Thus, inhibition of budding by hypoxia could prevent mitosis, leading to the G2 arrest.

**Figure 4:**
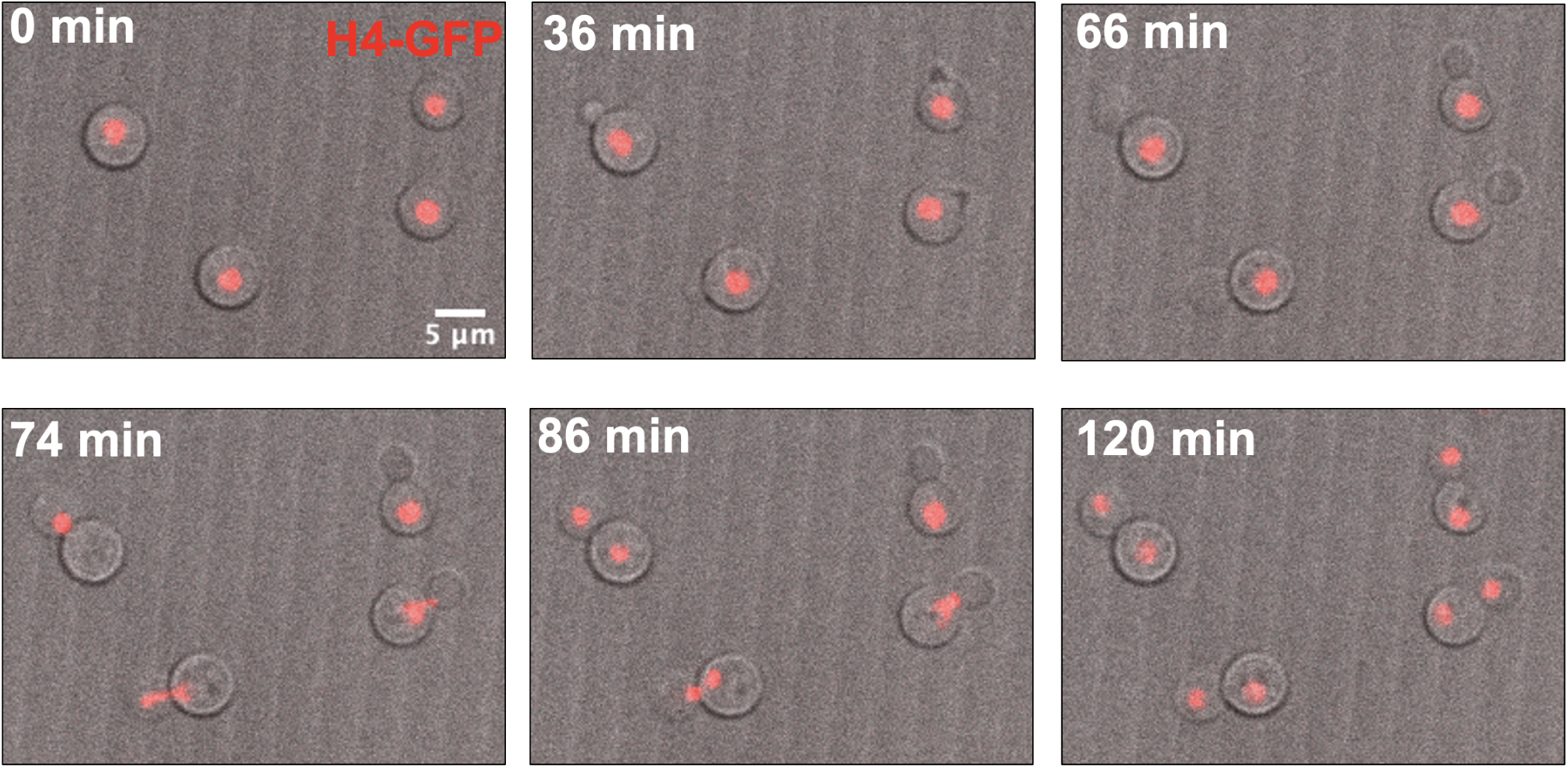
Budding precedes mitosis in *C. neoformans* cells after the release from saturation arrest. Live imaging of H99 cells released from saturation arrest onto fresh agarose pads (YEPD + 2% agarose). Chromatin is shown in red by GFP-tagged histone H4.

In *S. cerevisiae*, the morphogenesis checkpoint blocks mitotic progression in cells that fail to form a bud. In checkpoint mutants lacking the mitotic inhibitor Swe1, unbudded cells undergo mitosis in the mother cell, resulting in binucleate cells^16^. To test whether *C. neoformans* has a similar regulatory pathway, we treated *C. neoformans* cells with 10 µM Latrunculin A (Lat A) to block actin polymerization and budding. F-actin staining with phalloidin confirmed that actin structures were disrupted in Lat A-treated cells (Figure S5). If a morphogenesis checkpoint exists, cells treated with Lat A will arrest before mitosis in cells lacking a bud; otherwise, Lat A-treated cells will proceed into mitosis and become binucleated unbudded cells. Given that the *C. neoformans* cell cycle is approximately 70–80 min based on the synchronized budding curve (Figure S1), we treated *C. neoformans* cells for 90 and 120 min and *S. cerevisiae* cells for 90 min, thereby covering at least one full cell cycle such that any cells escaping checkpoint arrest would be expected to complete mitosis and become binucleate. As controls for morphogenesis checkpoint function, we treated *S. cerevisiae* wild-type cells and cells lacking the *SWE1* gene, which encodes a kinase required for morphogenesis checkpoint function^24,25^, with Lat A.

As expected, Lat A treatment inhibited budding in both *S. cerevisiae* and *C. neoformans* cells (Figure 5 A). Although Lat A treated cells accumulated a 2C DNA content consistent with G2/M arrest (Figure S6 A, B), unbudded cells did not produce a substantial number of binucleate cells in either species (Figure 5 B, C), suggesting that the block in budding prevented subsequent nuclear division. In contrast, *S. cerevisiae swe1*Δ cells produced binucleates in nearly 40% of cells despite Lat A treatment (Figure 5 B, C), consistent with the bypass of the morphogenesis checkpoint. These findings support the existence of a morphogenesis checkpoint in *C. neoformans*, and are consistent with a model in which hypoxia-induced inhibition of budding activates the morphogenesis checkpoint, leading to G2 arrest.

**Figure 5:**
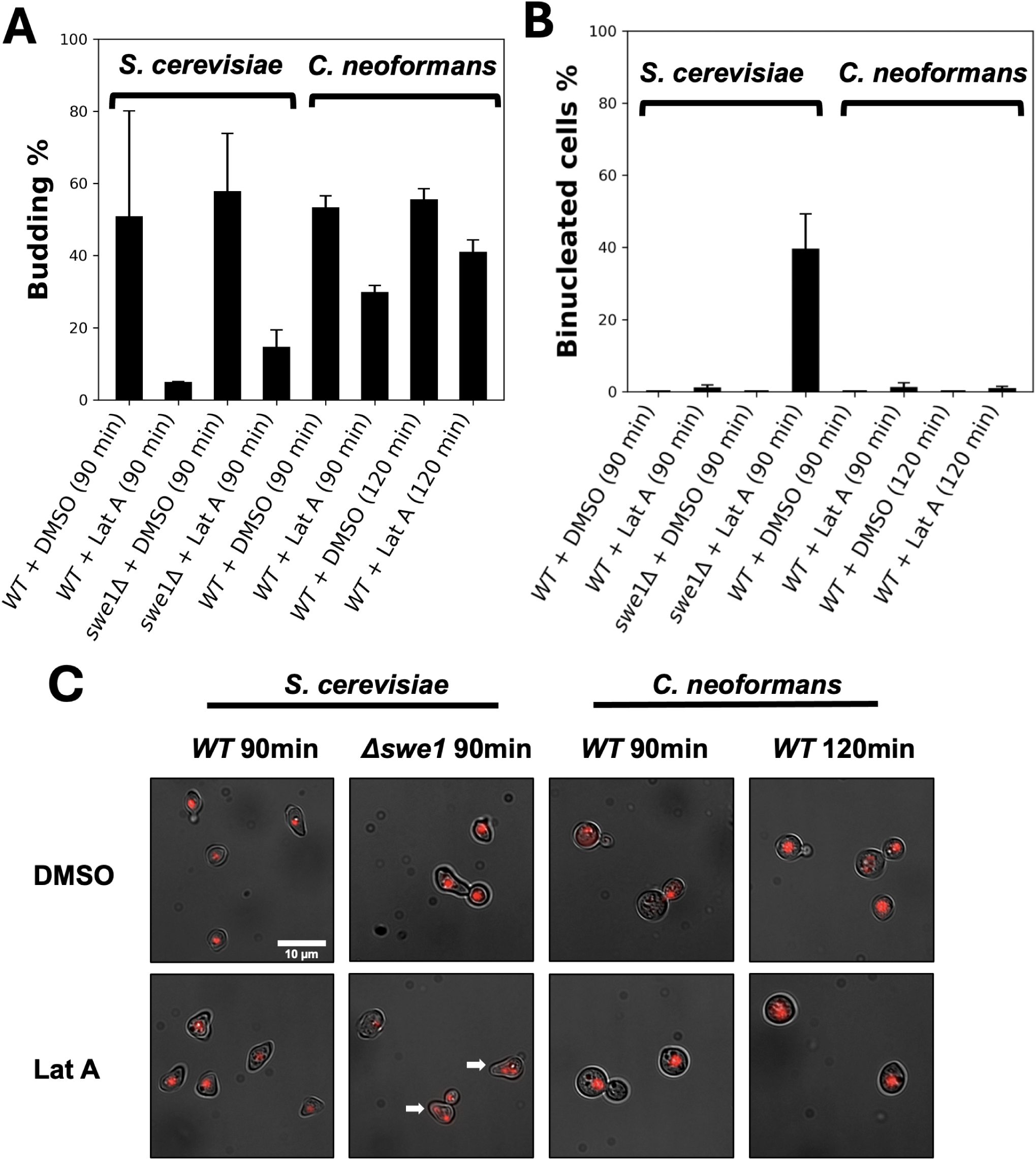
Mitosis is inhibited by Latrunculin A (Lat A) treatment in H99 cells. Percentages of budded cells (**A**), binucleated cells (**B**), and representative images (**C**) of *S. cerevisiae* wild-type (WT) and Δ*swe1* cells, and *C. neoformans* WT cells treated with DMSO or Lat A (final concentration is 10 µM). Red indicates DNA stained with SYTOX Orange. White arrows indicate binucleate cells.

### The morphogenesis checkpoint in *C. neoformans* is dependent on Swe1

To determine whether the morphogenesis checkpoint in *C. neoformans* was dependent on Swe1 as it is in *S. cerevisiae* ^25^, we repeated the Lat A experiment in *C. neoformans swe1*Δ mutant cells^26^ as well as wild-type cells (Figure 6 A,B,C and S7). Wild-type cells did not become binucleate when treated with Lat A; however, *swe1*Δ mutant cells exhibited more than 47 and 40 percent binucleates after 90 and 120 min, respectively. A small percentage of binucleate cells were also observed in the *swe1*Δ mutant control cells treated only with DMSO. Together, these findings indicated that a morphogenesis checkpoint in *C. neoformans* is enforced by Swe1, and that the checkpoint may be actively preventing the formation of binucleates even in normally dividing cells.

**Figure 6:**
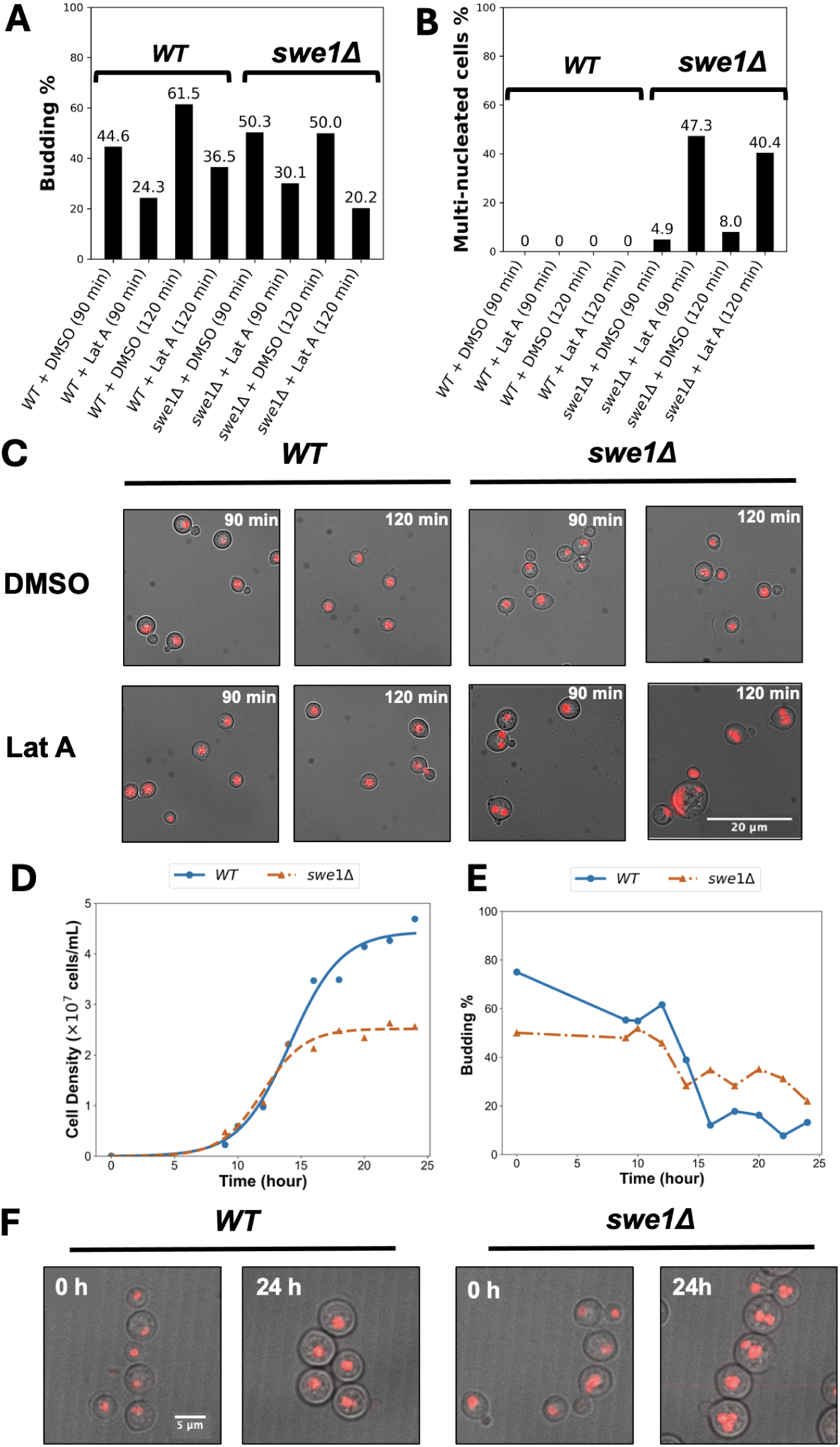
*C. neoformans swe1*; mutant becomes multinucleate through both Lat A treatment and saturation arrest. (**A**) Percentage of budded cells and (**B**) multinucleate cells in *C. neoformans* H99 wild type (WT) and *swe1*Δ mutant treated with DMSO or Lat A (10 µM). (**C**) Representative fluorescence images of the indicated strains and treatments. DNA is stained with SYTOX Orange (red). (**D**) Growth curves of H99 WT and *swe1*Δ fitted with a logistic model. (**E**) Budding index of the indicated strains over time. (**F**) Representative images of H99 WT and *swe1*Δ cells collected at 0 hour (early log phase) and 24 hour (saturation arrest). DNA is stained with SYTOX Orange (red).

Given that *C. neoformans* has a functional morphogenesis checkpoint that is activated when cells don’t bud, and that hypoxia appears to inhibit budding, the most straightforward explanation for the cell-cycle arrest in hypoxia is activation of the morphogenesis checkpoint. If true, then *C. neoformans swe1*Δ cells should not be mitotically arrested when they are subjected to hypoxia. Thus, we examined cell density, budding, and nuclear morphology in saturation arrested *swe1*Δ mutants (Figure 6 D, E, F).

Wild-type control cells were arrested as unbudded G2 cells at 24 hours. The *swe1*Δ cells saturated at a lower density than wild-type cells (Figure 6D) and were mostly mononucleated in early log phase (Figure S9). At 24 hours (approximately 10 hours after they arrested division), the *swe1*Δ mutant cells were largely polyploid (Figure S8), and many cells were multinucleated (Figure 6F and Video S2). Specifically, 36% of cells contained two nuclei and 23% contained three or more nuclei (Figure S9), indicating that nuclear division (mitosis) was not inhibited. This observation is consistent with a role for Swe1 in preventing mitosis in hypoxia when budding is inhibited. Although Swe1 could be the target of some other hypoxia-induced signaling pathway, the simplest explanation is that hypoxia is triggering the *C. neoformans* morphogenesis checkpoint.

## Discussion

It has been generally assumed that saturation-induced arrest in *C. neoformans* is caused by nutrient depletion^5^. Here, we show that instead, the arrest of cell division at saturation in liquid medium is caused by hypoxia. This finding is consistent with previous observations that low O_2_ levels were correlated with the unbudded G2 arrest in *Cryptococcus* ^19,27^. The previous studies also found that as cells start to become hypoxic, budding emergence appears to be shifted from early in the cell cycle to late in G2, and then inhibited completely as cells further deplete oxygen from the medium^20^(Figure 1). Although these reports suggested hypoxia was correlated with saturation arrest, the mechanism underlying hypoxia-induced G2 arrest remained unknown.

We discovered that inhibition of bud emergence in *C. neoformans* by treatment with Lat A arrested cells before mitosis via a Swe1-dependent morphogenesis checkpoint (Figures 5 and 6 A, B, C). We also demonstrated that a Swe1-dependent mechanism is responsible for arresting cells in response to hypoxia (Figure 6 D, E, F). Taken together, these findings suggest that hypoxia blocks budding and the lack of a bud triggers a morphogenesis checkpoint similar to that observed in *S. cerevisiae*. That said, our results do not rule out that Swe1 enforces some other hypoxia-induced checkpoint mechanism in addition to a morphogeneis checkpoint or that *C. neoformans* might directly monitor F-actin polymerization.

The mechanism by which hypoxia induces the shift in budding from early to late in the cell cycle and ultimately inhibits budding (Figure 1)^19,20,27^ remains unknown. It has been shown that the membrane-bound protein Sre1 serves as the oxygen sensor in *C. neoformans* ^28–30^. Under hypoxic conditions, Sre1 is cleaved by the protease Stp1, releasing its N-terminal portion into the cytosol. This cleaved fragment then functions as a transcription factor to activate genes required for ergosterol biosynthesis and iron uptake in response to hypoxia. Whether the Sre1 pathway is involved in the shift and inhibition of budding is a question for future investigations.

Aside from the involvement of Swe1, it also remains unclear whether the morphogenesis checkpoint mechanism in *C. neoformans* utilizes the same machinery observed in the *S. cerevisiae* morphogenesis checkpoint^16,24,31–33^. When unbudded G2 cells arrested by hypoxia are released in fresh oxygenated medium, the cells first form a bud before transporting the nucleus into the bud, where anaphase ultimately pushes one of the divided nuclei back into the mother cell (Figure 4; Kozubowski et al^23^). This observation contrasts with anaphase dynamics observed in *S. cerevisiae*, where the nucleus is positioned in the mother cell at the bud neck, and the anaphase spindle moves one of the divided nuclei into the bud^34^. These findings suggest the morphogenesis checkpoint in *C. neoformans* might monitor the completion of nuclear migration into the bud rather than bud emergence itself. Under this model, entry into anaphase may require nuclear migration into the bud because the bud compartment provides signals that permit anaphase onset. How nuclear migration at mitosis is controlled in *C. neoformans* is also a topic for future studies.

Cell-cycle checkpoints are reversible intracellular signaling pathways that monitor cell-cycle progression and ensure that cell-cycle events are executed in the proper order^35^. Checkpoint pathways evolved to sense a perturbation (e.g. DNA damage) and then enable the cell to actively pause cell division before genome stability or cell viability are irreversibly impacted^12,35^. For example, the DNA replication checkpoint detects DNA damage and stops the cell cycle in S phase, allowing time for repair. If mutations in checkpoint genes disrupt the arrest, cells continue cell-cycle progression despite DNA damage, leading to rapid loss of viability, as observed in *S. cerevisiae rad9*Δ cells with damaged DNA.^36,37^. Once the perturbation is resolved (e.g. repair of DNA damage) checkpoint signaling ends, allowing the cells to reverse the arrest and re-initiate cell division. Thus, checkpoints are active mechanisms that protect genome integrity and cell viability from environmental stresses.

*Cryptococcus* species are obligate aerobes^38,39^, hypoxia is known to trigger global metabolic reprogramming in *C. neoformans*, including changes in energy metabolism, mitochondrial function, and sterol biosynthesis^28,29,40,41^, all of which are important for cell division and survival. Thus, the hypoxia-induced G2 checkpoint we have identified likely evolved to protect cell viability under hypoxic stress. Here we show that cells lacking this checkpoint (*swe1*Δ mutant cells) became extensively multinucleated and polyploid under hypoxic conditions (Figure 6, S8 and Video S2). Polyploidization and abnormal nuclear division are frequently associated with genome instability and can impair cell fitness^42^. Therefore, activation of the checkpoint allows cells to exit the cell cycle in G2 before cells become multinucleated, and protects the cells from becoming polyploid in hypoxia, preserving genome integrity, and potentially improving cell viability.

Our findings may have important implications for understanding the mechanism of Titan cell formation in *Cryptococcus*. G2-unbudded cells have been reported as precursors to Titan cells^4^, and here we show that an unbudded G2 arrest can be induced by hypoxia. One of the key properties of Titan cells is polyploidy, which is thought to be generated through endocycling, or endoreplication mechanisms^43–45^. Endocycling requires cells to block mitosis while continuing repeated cycles of G and S phases. The hypoxia-induced G2 arrest we observed could provide precisely such a mechanism to block mitosis, offering a possible trigger for the initiation of endocycling. Polyploidy could arise from this G2 arrest if the checkpoint we identified is not fully capable of maintaining a cell cycle arrest over time, as suggested by the DNA content of the *swe1*Δ cells (Figures S7 and S8), which eventually became polyploid following prolonged arrest. In fact, checkpoints are commonly reported to be “leaky” and allow cells to re-enter the cell cycle after prolonged arrest^46–48^. Future studies will focus on investigating potential connections between the hypoxia pathway and titan cell formation.

## RESOURCE AVAILABILITY

### Lead contact

Requests for further information and resources should be directed to and will be fulfilled by the lead contact, Steven B. Haase.

### Materials availability

Strains in this study will be made available upon request.

### Data and code availability

- All data reported in this paper will be shared by the lead contact upon request.
- This paper does not report original code.
- Any additional information required to reanalyze the data reported in this paper is available from the lead contact upon request.

## ACKNOWLEDGMENTS

This work was funded by the NIH **1R21AI193612-01**. Many thanks to Esther Miranda and David R. McClay for offering an air pump to this study and to Daniel P. Kiehart for help in the oxygen measurement. We would also thank Duke Light Microscopy Core Facility and the director, Lisa Cameron, for help in microscopy. *C. neoformans swe1*Δ strains were a gift from Yong-Sun Bahn and Lukasz Kozubowski. We thank Daniel Lew, Joseph Heitman, Andrew Alspaugh, Vikas Yadav, Yong-Sun Bahn, and Lukasz Kozubowski for helpful discussions and critical review of the manuscript.

## AUTHOR CONTRIBUTIONS

Conceptualization, H.Z. and S.B.H.; methodology, H.Z., A.H.L., and C.P.; investigation, H.Z., A.H.L., and C.P.; writing-–original draft, H.Z. and S.B.H.; writing-–review & editing, H.Z., A.H.L., C.P. and S.B.H.; funding acquisition, S.B.H.; supervision, S.B.H.

## SUPPLEMENTAL INFORMATION INDEX

Figures S1–S8 and their legends are provided at the end of the manuscript. Video S1 and Video S2 provided as separate MP4 files.

- Video S1. Cells released from hypoxia arrest. Cells form buds, nuclei move into buds, then mitosis occurs. Maximum projection merged time series of GFP-H4 at 2-minute intervals. The playback speed is 5 frames per second.
- Video S2. Rotating 3D projection of SYTOX Orange-stained *swe1*Δ *C. neoformans* cells at 24 hour saturation arrest, revealing multinucleate morphology. Scalebar = 2 *µ*m.

## STAR METHODS

### Key resources table

| REAGENT or RESOURCE | SOURCE | IDENTIFIER |
| --- | --- | --- |
| **Chemicals** |  |  |
| 2-mercaptoethanol | Bio-Rad | Cat. no. 1610710 |
| Sodium ascorbate | Nutricost | Cat. no. NCS-57184 |
| RNase A | Qiagen | Cat. no. 19101 |
| Dimethyl sulfoxide (DMSO) | Sigma-Aldrich | Cat. no. D8418 |
| Formaldehyde solution | Sigma-Aldrich | Cat. no. F8775 |
| Latrunculin A | Sigma-Aldrich | Cat. no. 428026 |
| Paraformaldehyde (PFA) | Sigma-Aldrich | Cat. no. 158127 |
| Sodium hydroxide | Sigma-Aldrich | Cat. no. 901915 |
| α1-Mating Factor acetate salt | Sigma-Aldrich | Cat. no. T6901 |
| Alexa Fluor™ 488 Phalloidin | Thermo Fisher Scientific | Cat. no. A12379 |
| SYTOX™ Orange Nucleic Acid Stain | Thermo Fisher Scientific | Cat. no. S11368 |
| **Software and Algorithms** |  |  |
| FlowCal | Castillo-Hair et al., 2016<sup>49</sup> | https://flowcal.readthedocs.io |
| Python | Python Software Foundation | Version 3.10.10 |
| Matplotlib | Matplotlib Development Team | Version 3.5.2 |
| NumPy | Harris et al., 2020<sup>50</sup> | Version 1.24.3 |
| SciPy | Virtanen et al., 2020<sup>51</sup> | Version 1.11.1 |
| FIJI (ImageJ) | Schindelin et al., 2015<sup>52</sup> | https://imagej.net/Fiji |
| MetaMorph | Molecular Devices | N/A |
| SoftWoRx | Applied Precision | Version 6.1 |
| **Instruments** |  |  |
| QSonica Sonicator Q55 | QSonica | N/A |
| Attune NxT Flow Cytometer | Thermo Fisher Scientific | N/A |
| Opto-F4 oxygen meter | UniAmp | N/A |
| Andor Revolution XD microscope | Andor Technology | N/A |
| DeltaVision Elite microscope | Applied Precision | N/A |

### Experimental model and study participant details

*Saccharomyces cerevisiae* strains used in this study were in the 15D MATa background. The 15D MATa strain was used as the wild type. The *swe1*Δ strain (*bar1, swe1::URA3 GAL1-CDC28::TRP1 pGAL-CLB2-TAP::URA3*; DLY5806) was a gift from Daniel Lew.

*Cryptococcus neoformans* strains used in this study were in the H99 background (H99S). The H99 strain was used as the wild type. The GFP-H4 strain (*GFP-H4::NAT*; VYD158) was a gift from Joseph Heitman and described previously^23^. The *swe1*Δ strains (*swe1*Δ::NAT; YSB8309, YSB8311, YSB8312)^26^ were gifts from Yong-Sun Bahn and Lukasz Kozubowski. The *swe1*Δ strains were verified by PCR using primers

- swe1 Fw (5’ ACTTACCCTCCCTGGATTGACGTG 3’)
- swe1 Rv (5’ CTGGTGTAATCAAGCCCATCCCTC 3’)
- NAT middle Rv (5’ GTTCCAGCCGGAGTACGAGACG 3’).

swe1 fw and swe1 Rv flank the *SWE1* locus, whereas NAT middle Rv binds within the NAT resistance cassette.

*Cryptococcus gattii* R265 and *Cryptococcus deneoformans* JEC21 strains were gifts from Joseph Heitman.

Strains were cultured at 30 °C in YEPD liquid medium (1% [w/v] yeast extract, 2% [w/v] peptone, 2% [w/v] dextrose, 0.012% [w/v] adenine hemisulfate, and 0.006% [w/v] uracil) or on YEPD agar plates containing 1.5% (w/v) agar. Strains were stored at −80 °C in a mixture of 1 mL yeast culture grown in YEPD medium and 1 mL of 50% glycerol solution.

### Method details

#### Generation of saturation-arrested cells

*C. neoformans* cells were inoculated into 100 mL YEPD medium in a 250 mL glass Erlenmeyer flask with a cap at a starting density of approximately 2 × 10^6^ cells/mL. Cells were cultured at 30 °C in a shaking water bath at 120 rpm.

For cell counting, 200 µL of culture was mixed with 200 µL of 4% paraformaldehyde (PFA), and cells were counted using a hemocytometer under a light microscope. Budding percentage was defined as the proportion of budded cells among the total cell population.

After 24 hours of growth, cultures reached saturation and consisted G2 cells (saturation arrest).

#### Growth curve fitting

Cell density measurements were fitted with a three-parameter logistic model of the form:

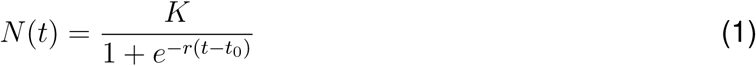

where *K* is the carrying capacity, *r* is the growth rate (h*^−^*^1^), and *t*_0_ is the inflection point (h). Parameters were estimated by nonlinear least-squares regression using SciPy (v1.11.1;^51^) in Python. All data visualization was performed using Matplotlib.

#### Flow cytometric analysis of DNA content

Flow cytometry was used to assess cellular DNA content. A total of 1–5 × 10^7^ cells were collected by centrifugation at 3,000 × g for 4 min, washed once with 3 mL deionized water, and resuspended in 0.5 mL of water. Subsequently, 1.3 mL of 100% ethanol was added slowly with gentle mixing to obtain a final concentration of 70%(v/v) ethanol. Cells were fixed at room temperature for at least 1 hour and then stored at 4 °C overnight or longer.

After fixation, cells were pelleted by centrifugation (3,000 × g for 4 min), washed once with deionized water, and resuspended in 500 µL 50 mM Tris-HCl (pH 7.5). A 30 µL aliquot of the cell suspension was mixed with 500 µL staining solution (1 µM SYTOX Orange [cat. no. S11368; Thermo Fisher Scientific] in 50 mM Tris-HCl (pH 7.5), 2.5 µL RNase A [10 mg/mL; cat. no. 19101; Qiagen], and 1 µL 2-mercaptoethanol [cat. no. 1610710; Bio-Rad]). Samples were incubated overnight at 4 °C in the dark.

Immediately prior to analysis, samples were briefly sonicated (QSonica Sonicator Q55, amplitude:15, duration:10 s) to disperse cell aggregates. Flow cytometry was performed using an Attune NxT Flow Cytometer (Thermo Fisher Scientific). Forward scatter (FSC) and the YL1 fluorescence channel were used for analysis. Instrument settings were as follows: FSC voltage 100, YL1 voltage 280, FSC threshold 5,000, and YL1 threshold 2,000. The acquisition volume was set to 40 µL with a flow rate of 25 µL/min, and data collection was stopped after 10,000 events.

Flow cytometry data were analyzed using Python (version 3.10.10) with FlowCal (version 1.3.0)^49^, and plots were generated using Matplotlib (version 3.5.2).

#### Oxygen level measurement

Dissolved oxygen levels in the medium were measured using an Opto-F4 oxygen meter (Uni-Amp). For calibration, deionized water bubbled with air for 5 min was used as the 100% air-saturated solution. A zero-oxygen calibration solution was prepared by dissolving 2 g sodium ascorbate (Nutricost) and 0.4 g sodium hydroxide (Sigma-Aldrich) in 100 mL deionized water. A linear calibration curve was generated based on the measured Δ*ϕ* values in 100% air-saturated solution and the zero-oxygen calibration solution.

During measurements for liquid culture, the probe was placed approximately 3 cm below the medium surface. After the Δ*ϕ* reading stabilized, the value was recorded and converted to dissolved oxygen saturation percentage according to the calibration curve.

#### Generation of hypoxic medium

Hypoxic medium was generated using two methods.

For the first method, YEPD medium was heated to boiling in a microwave and immediately transferred into a 150 mL glass flask (completely filled) containing a magnetic stir bar. The flask was capped, sealed with tape, and cooled at 4 °C. Before experiments, the medium was prewarmed at 30 °C. After inoculation, the flask was immediately sealed and placed on a magnetic stirrer to maintain mixing during growth. The flask was resealed immediately after each sampling.

For the second method, nitrogen gas was used to displace oxygen. A 150 mL flask was filled completely with medium. Liquid nitrogen was placed in a foam container and covered with a lid. Nitrogen gas evaporating from the liquid nitrogen was pumped into the medium while dissolved oxygen levels were monitored. Bubbling was continued until oxygen levels stabilized. The flask was then sealed with tape. Subsequent culture conditions were the same as described above.

#### Latrunculin A treatment

To inhibit budding using Latrunculin A (Lat A), cells were first synchronized to obtain predominantly unbudded populations.

For *Saccharomyces cerevisiae* wild-type and *swe1*Δ cells, log-phase cultures were treated with 15 ng/mL *α*-factor to arrest cells at the unbudded G1/S phase.

For *Cryptococcus neoformans* H99 wild-type cells, cultures were first grown to saturation arrest (100 mL culture, 24 h; see above) to synchronize cells in an unbudded state. Cells were then released into fresh medium and allowed to grow for 85 min until after the first division, when budding percentage reached its minimum (Figure S1). Cells were then collected by centrifugation at 1,500 × g for 4 min, the medium was removed, and cells were kept on ice.

For *C. neoformans* H99 *swe1*Δ cells, saturation arrest resulted in multinucleated cells that interfered with analysis. Therefore, cultures were grown for 16 hours instead of 24 hours until the budding percentage decreased, followed by release into fresh medium for 85 min before centrifugation.

After obtaining mostly unbudded cells, cells were resuspended in 500 µL YEPD medium in glass tubes. Either 5 µL DMSO (cat. no. D8418; Sigma-Aldrich) or 5 µL Latrunculin A (1 mM stock; cat. no. 428026; Sigma-Aldrich) was added. The final Lat A concentration was 10 µM. Cultures were mixed and incubated at 30 °C on a rotating platform.

#### Phalloidin staining

Cells were collected and pelleted by centrifugation at 1,500 × *g* for 4 min. The supernatant was removed and cells were resuspended in 1 µL fixation solution (4% formaldehyde; 100 µL 36% formaldehyde [Sigma-Aldrich, F8775] diluted in 900 µL PBS) and incubated at room temperature for 30 min. Cells were then pelleted at 12,000 rpm for 1 min, the supernatant was removed, and cells were resuspended in 100 µL phalloidin staining solution (2 U/mL Alexa Fluor™ 488 Phalloidin [Thermo Fisher Scientific, A12379], 0.5% Triton X-100, 1× PBS) and incubated at 4 °C on a shaker overnight (about 16 hours). Prior to imaging, cells were pelleted at 12,000 rpm for 1 min and resuspended in 50 µL PBS at 4 °C.

#### Cell imaging

To grow cells for imaging of release from hypoxia arrest, cells were cultured for 24 hour at 30°C in YEPD with shaking at 110 rpm. After 24 hour, most cells were arrested as unbudded in G2. Cells were sonicated for 4 s, and at this time we assumed cells were released from hypoxia arrest. 3 µL of cells were placed in an ibidi 8-well glass-chamber µ-slide with a 200 µL slab of YEPD medium on top, solidified with 2% agarose.

To observe the order of bud emergence and mitosis, time-lapse images of cells expressing H4-GFP were acquired using an Andor Revolution XD spinning disk confocal microscope (Andor Technology) equipped with a CsuX-1 5,000 rpm confocal scanner unit (Yokogawa) and a UPLSAPO 60×/1.3 silicone oil objective (Olympus) controlled by MetaMorph software (Molecular Devices). Images were captured by an iXon Life 888 EMCCD camera (Andor Technology). The entire cell volume was acquired using 14 Z-slices at 0.48 µm steps. For imaging the GFP-H4 probe, we used exposure time of 250 ms at 10% laser power (excitation 488 nm). DIC images were acquired with an exposure time of 50 ms. Images were acquired at 2 min intervals.

To image nuclei in wild-type and *swe1*Δ cells in hypoxia, cells were fixed and stained with SYTOX Orange (same as for flow cytometry, see above). Cells were mounted on slabs containing water solidified with 2% agarose [Bio-rad]. Images were acquired using an Andor Revolution XD spinning disk confocal microscope (Andor Technology) equipped with a CsuX-1 5,000 rpm confocal scanner unit (Yokogawa) and a UPLSAPO 100×/1.4 oil objective (Olympus) controlled by MetaMorph software (Molecular Devices). Images were captured by an iXon Life 888 EMCCD camera (Andor Technology). The entire cell volume was acquired using Z-slices at 0.3 µm steps. To image SYTOX orange, we used exposure time of 200 ms at 3% laser power (excitation 561 nm). DIC images were acquired with an exposure time of 50 ms.

Lat A-treated cells were imaged using a DeltaVision Elite microscope (Applied Precision) equipped with an Olympus IX-71 base and a 60×/1.42 NA oil immersion objective (PLAPON60XO Images were acquired using a CoolSNAP HQ2 CCD camera with SoftWoRx 6.1 software. SYTOX Orange-stained nuclei were imaged using the mCherry/AF594 channel (excitation 575/25 nm, emission 632/60 nm) at 50% illumination intensity with an exposure time of 0.2 s. Alexa Fluor™ 488 Phalloidin staining was imaged using the GFP/FITC channel (excitation 475/28 nm, emission 525/50 nm) at 50% illumination intensity with an exposure time of 0.2 s. Differential interference contrast (DIC) images were acquired using the polarization channel at 10% illumination intensity with an exposure time of 0.05 s.

All image analysis was performed in FIJI^52^.

3D projections of SYTOX stained wild type and *swe1*Δ cells were generated in FIJI using the 3D projection tool with the brightest-point projection method, rotation around the *y*-axis, and a slice spacing of 0.3 µm to match the *z*-step size used during image acquisition. Interpolation was enabled for all projections. Playback speed is 7 frames per second.

## 1 SUPPLEMENTAL FIGURES

**Figure S1:**
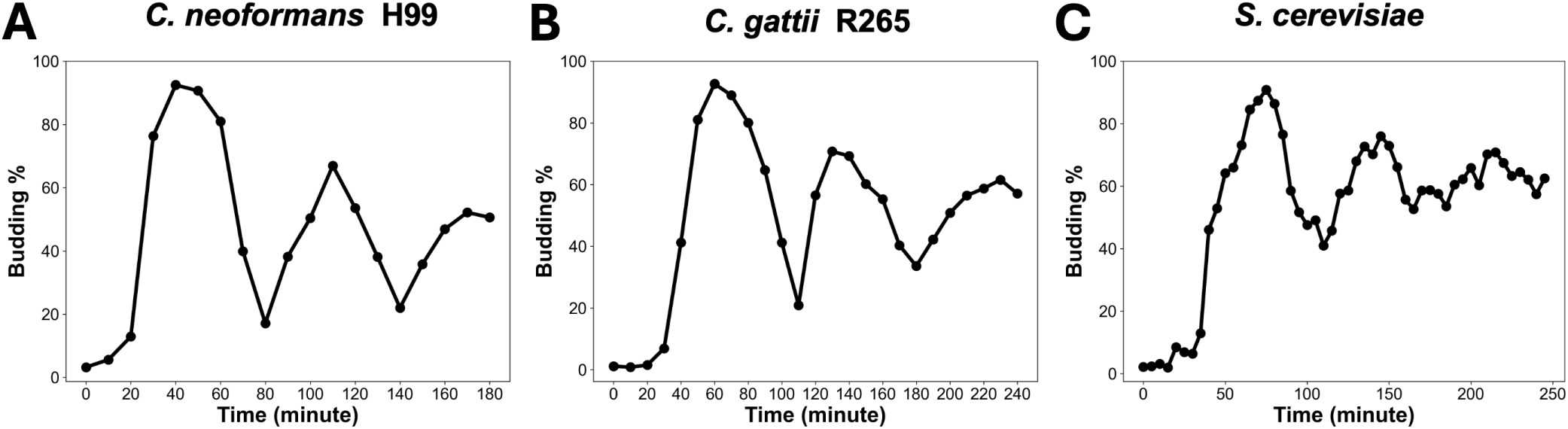
Budding curves of *C. neoformans* H99 (**A**) and *C. gattii* R265 (**B**) synchronized by saturation arrest, and *S. cerevisiae* (**C**) synchronized by alpha factor.

**Figure S2:**
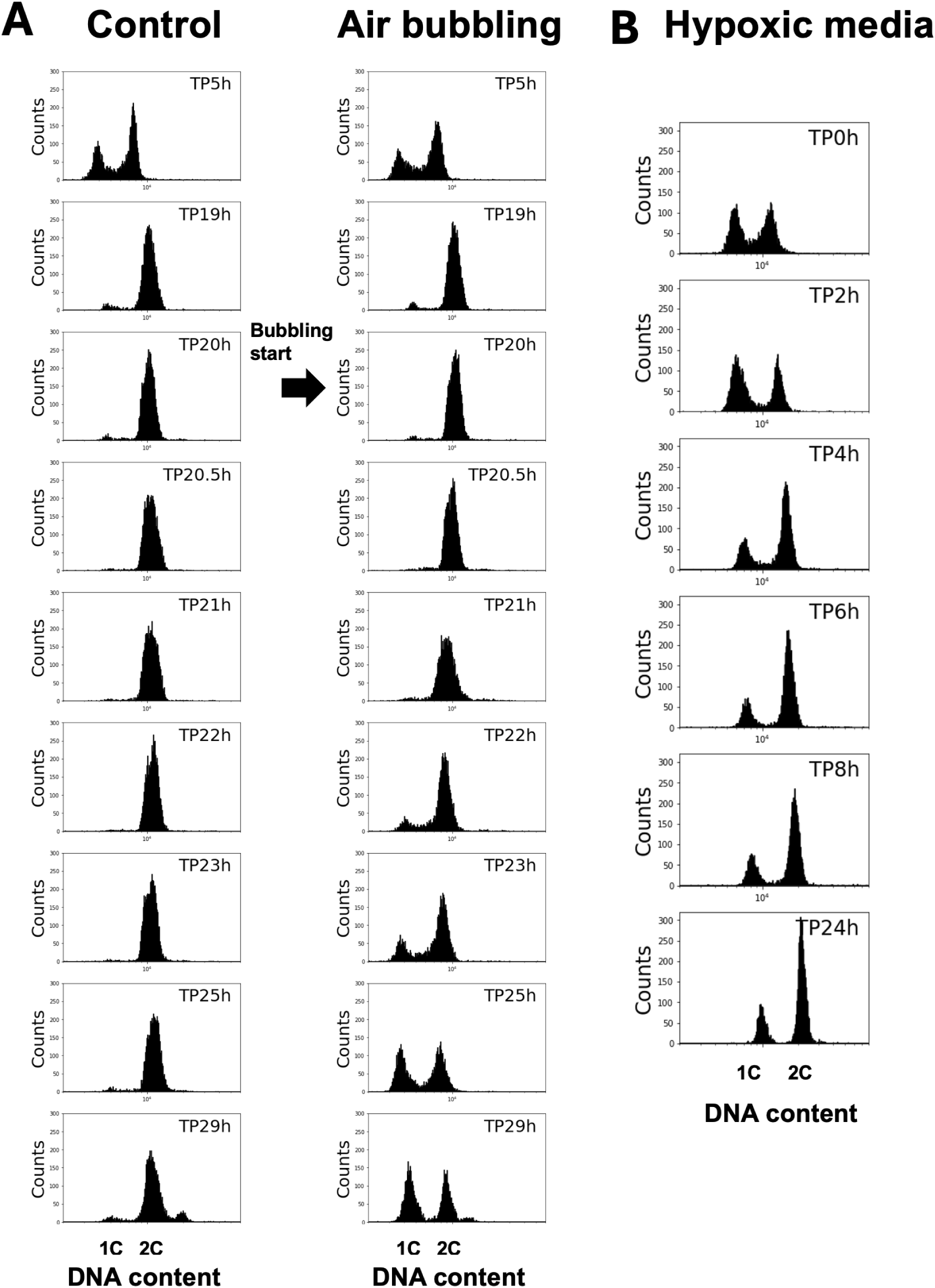
DNA content of cells shown in Figure 3, including H99 cells grown without air bubbling (**A**), with air bubbling initiated at the time indicated by the arrow (**B**), or in hypoxic medium generated by boiling (**C**). DNA was stained with SYTOX Orange.

**Figure S3:**
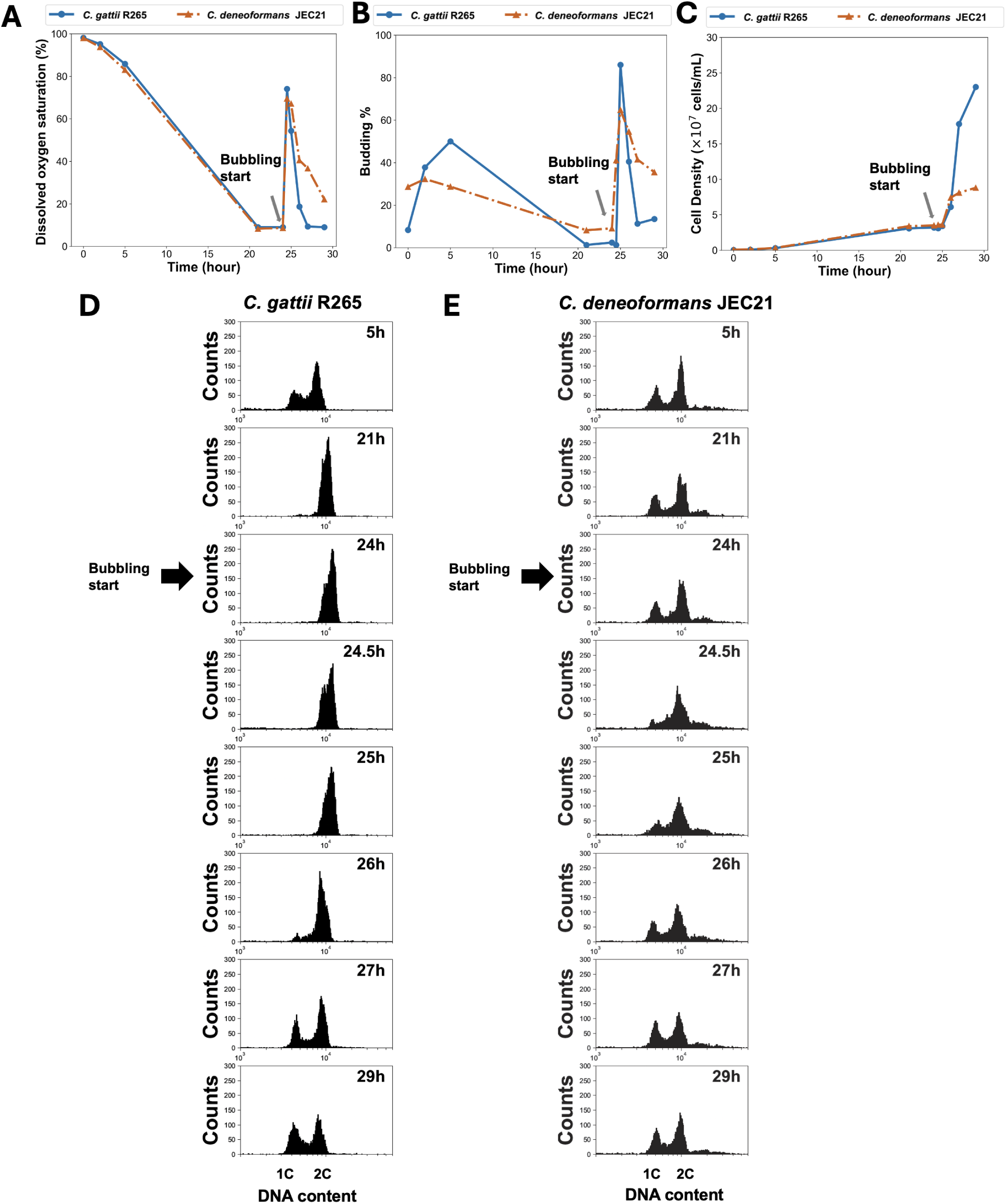
Oxygen level (**A**), budding index (**B**), and growth curves (**C**) of *C. gattii* R265 and *C. deneoformans* JEC21 cells with air bubbling initiated at the time indicated by the arrow. DNA content of cells collected from the same experiments: R265 (**D**) and JEC21 (**E**). DNA was stained with SYTOX Orange.

**Figure S4:**
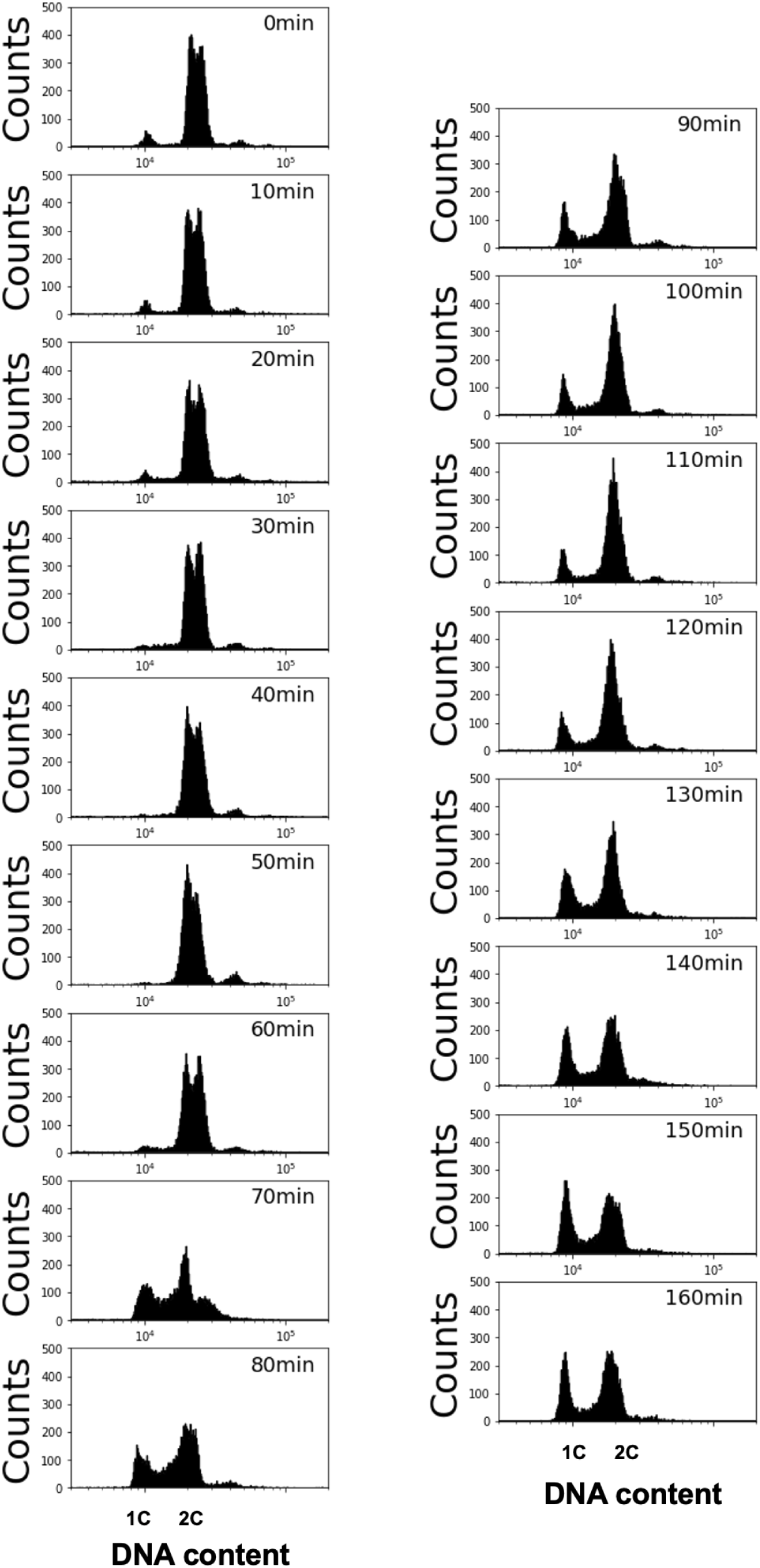
DNA content of the same cell population (as in Figure 4) released from saturation arrest into fresh medium and grown in parallel in liquid culture during live imaging. DNA was stained with SYTOX Orange.

**Figure S5:**
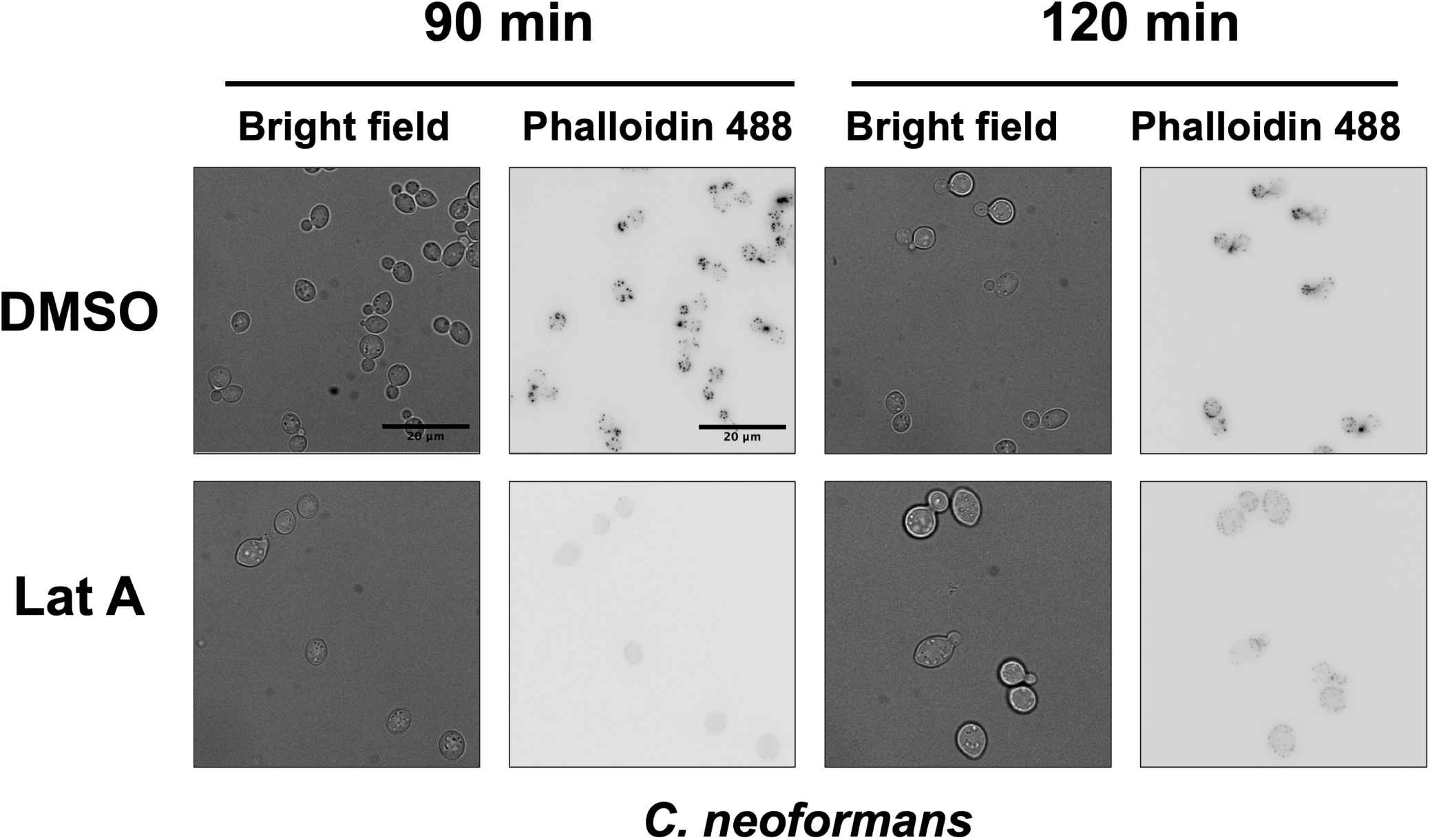
Lat A disrupts actin in H99 cells. Phalloidin 488 staining of *C. neoformans* H99 cells treated with DMSO (control) or Latrunculin A (Lat A) for 90 or 120 minutes.

**Figure S6:**
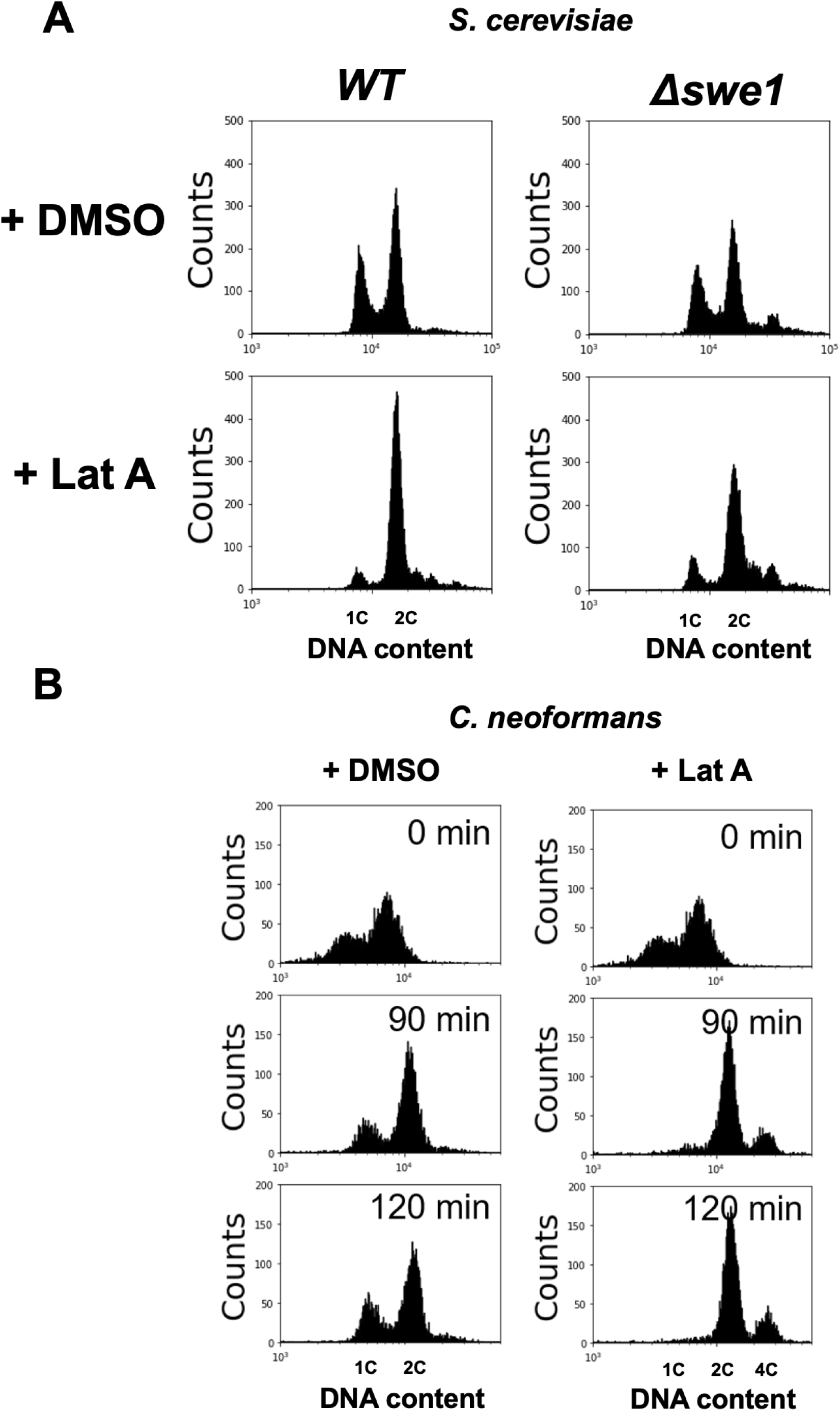
DNA content of cells treated with DMSO (control) or the actin inhibitor Latrunculin A (Lat A). (**A**) *S. cerevisiae* wild-type (15Da) and *swe1*Δ strains treated for 90 min. (**B**) *C. neoformans* H99 treated for 0, 90, and 120 min. DNA was stained with SYTOX Orange.

**Figure S7:**
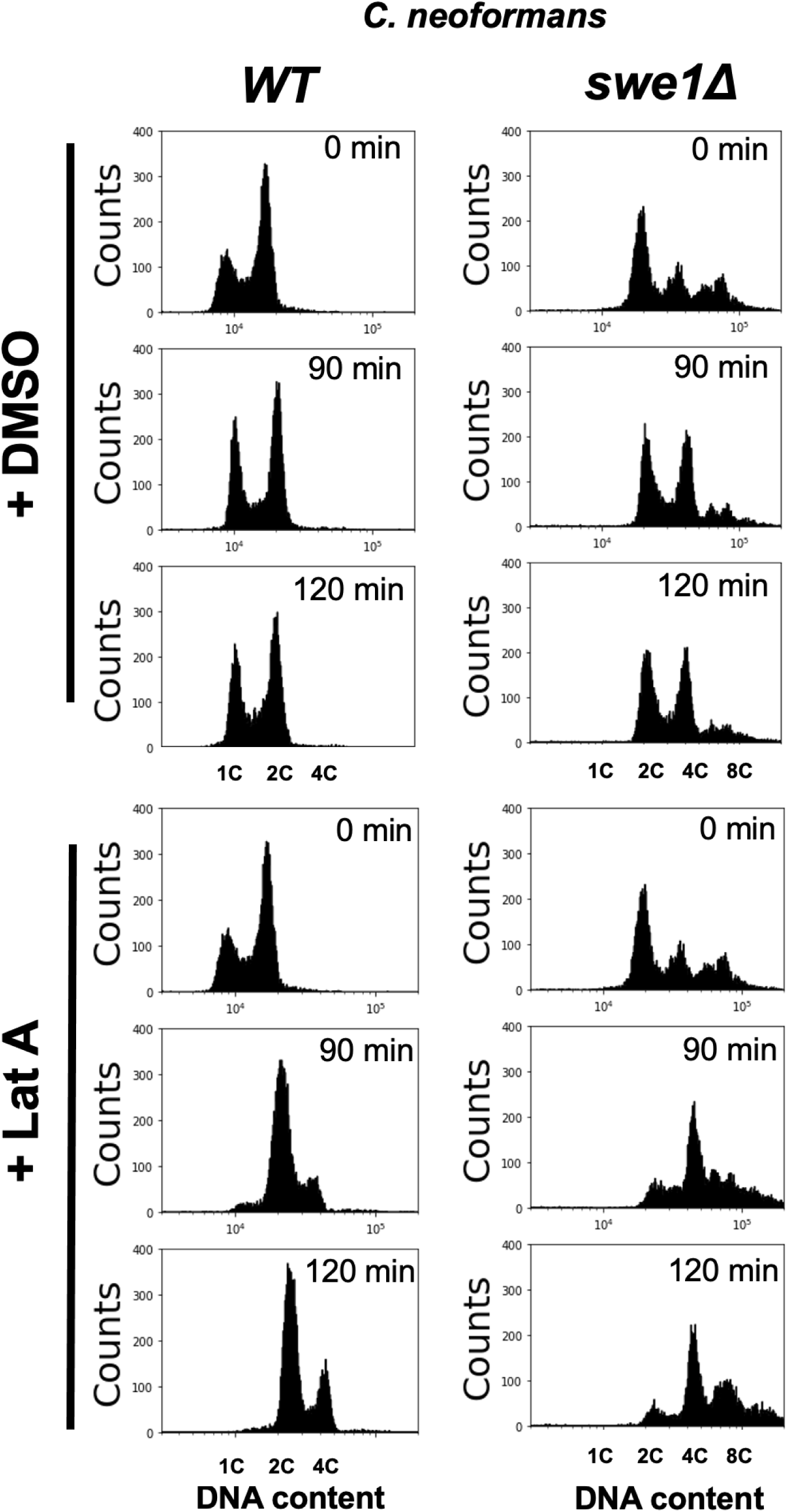
DNA content of cells shown in Figure 6 A,B,C. DNA was stained with SYTOX Orange.

**Figure S8:**
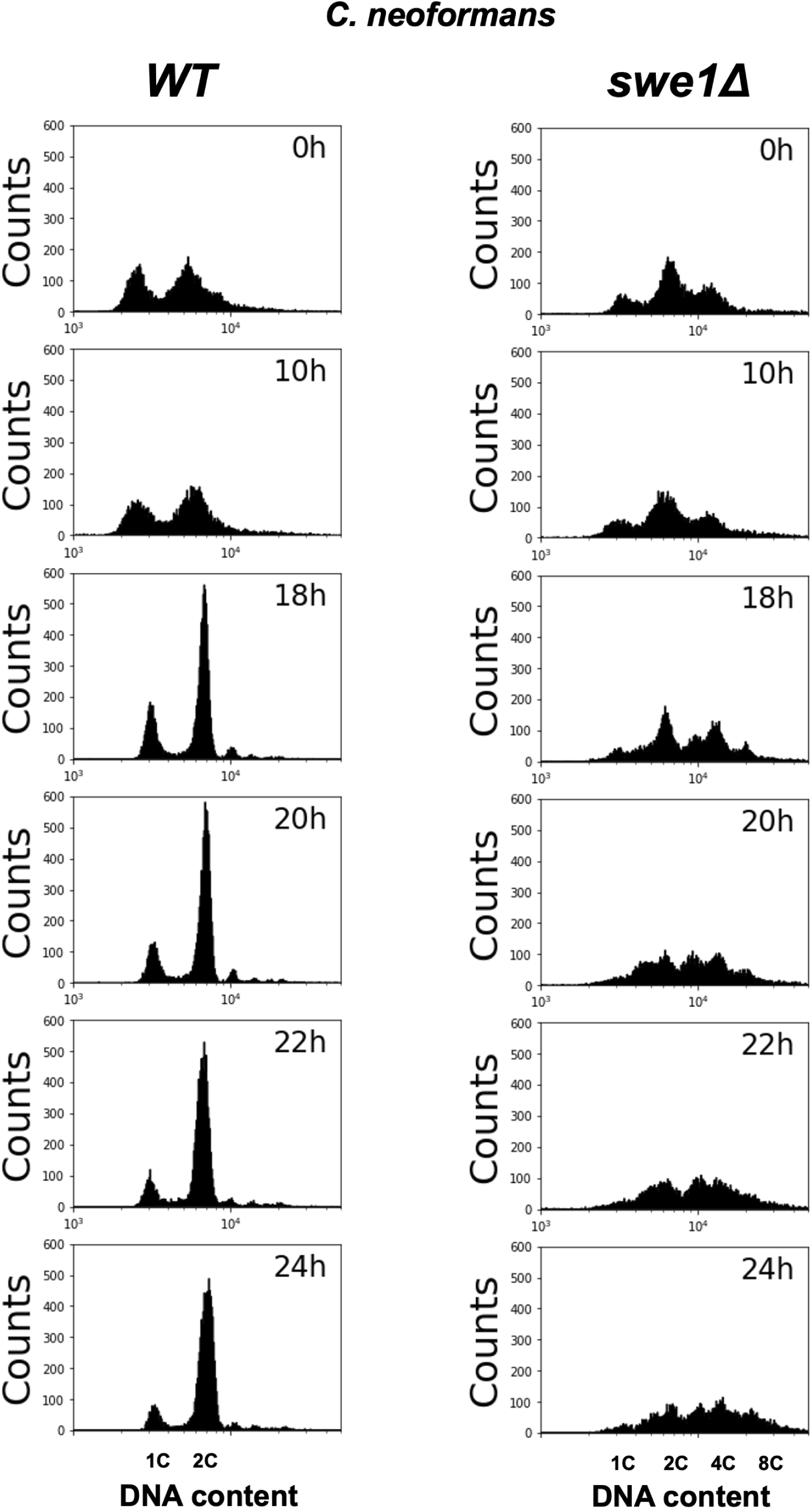
DNA content of cells shown in Figure 6 D,E,F. DNA was stained with SYTOX Orange.

**Figure S9:**
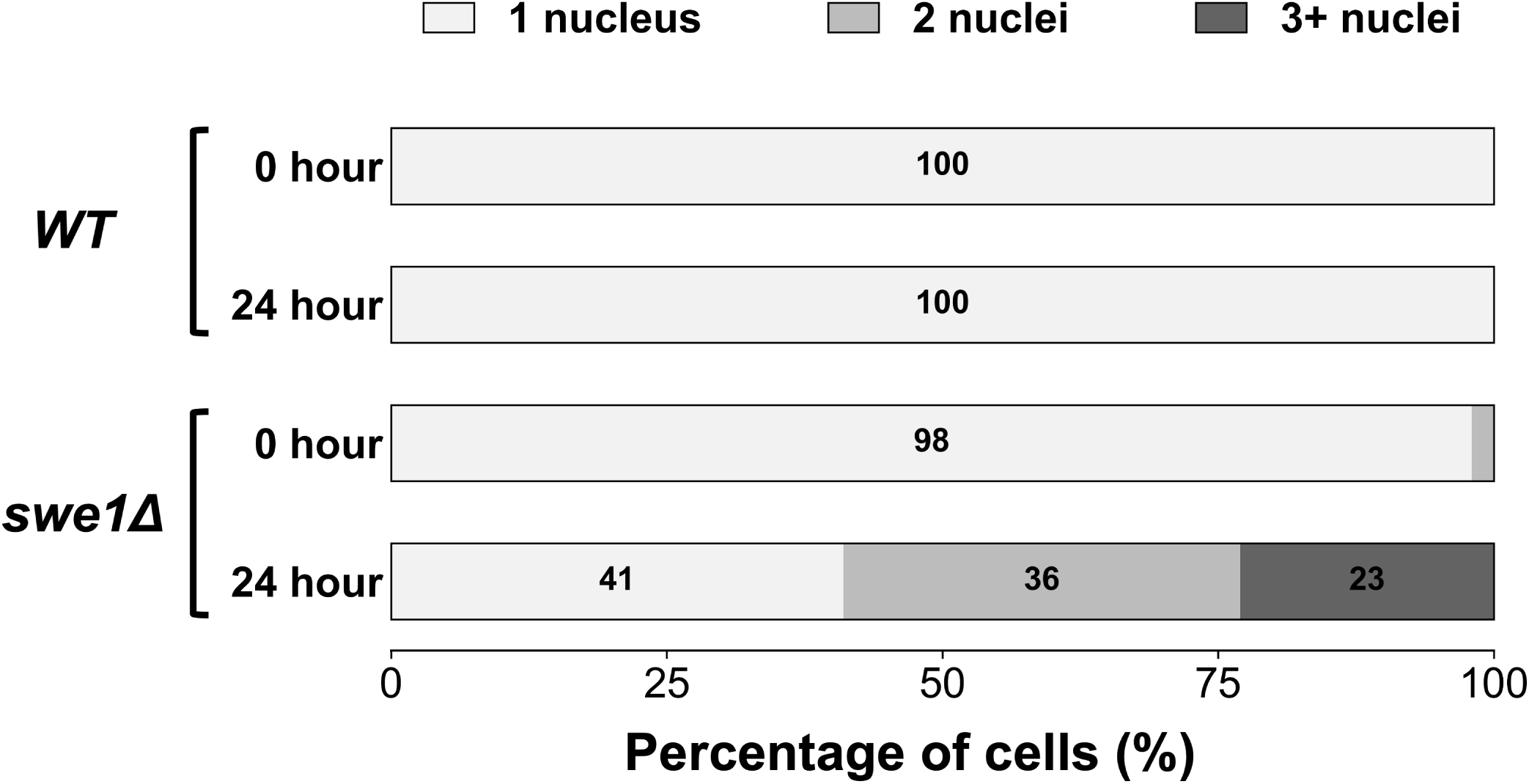
Percentage of unbudded cells containing one, two, or three or more nuclei in *C. neoformans* wild-type and *swe1*Δ strains shown in Figure 6 F. Only unbudded cells were counted to exclude potential interference from budded cells, in which two nuclei may occur as part of normal mitosis.

